# Comparative Phenotyping of Two Commonly Used *Chlamydomonas reinhardtii* Background Strains: CC-1690 (*21gr*) and CC-5325 (the CLiP Mutant Library Background)

**DOI:** 10.1101/2022.01.15.476454

**Authors:** Ningning Zhang, Leila Pazouki, Huong Nguyen, Sigrid Jacobshagen, Ming Xia, Anastasiya Klebanovych, Kirk J. Czymmek, Ru Zhang

**Author notes:** Corresponding author: Ru Zhang. Equal contribution (Ningning Zhang, Leila Pazouki, Huong Nguyen).

## Abstract

The unicellular green alga *Chlamydomonas reinhardtii* is an excellent model organism to investigate many essential cellular processes in photosynthetic eukaryotes. Two commonly used background strains of *Chlamydomonas* are CC-1690 and CC-5325. CC-1690, also called *21gr*, has been used for the Chlamydomonas genome project and several transcriptome analyses. CC-5325 is the background strain for the *Chlamydomonas* Library Project (CLiP). Photosynthetic performance in CC-5325 has not been evaluated in comparison with CC-1690. Additionally, CC-5325 is often considered to be cell-wall deficient, although detailed analysis is missing. The circadian rhythms in CC-5325 are also unclear. To fill these knowledge gaps and facilitate the use of the CLiP mutant library for various screens, we performed phenotypic comparisons between CC-1690 and CC-5325. Our results showed that CC-5325 grew faster heterotrophically in dark and equally well in mixotrophic liquid medium as compared to CC-1690. CC-5325 had lower photosynthetic efficiency and was more sensitive to heat than CC-1690. Furthermore, CC-5325 had an intact cell wall with comparable integrity to that in CC-1690, though appears to have reduced thickness. Finally, CC-5325 could perform phototaxis, but could not maintain a sustained circadian rhythm of phototaxis as CC160 did. Our results will be useful for researchers in the *Chlamydomonas* community to choose suitable background strains for mutant analysis and employ the CLiP mutant library for genome-wide mutant screens under appropriate conditions, especially in the areas of photosynthesis, thermotolerance, cell wall, and circadian rhythms.

## Introduction

The unicellular green alga *Chlamydomonas reinhardtii* (Chlamydomonas throughout) is a superior model organism to study important cellular processes in photosynthetic eukaryotes, including but not limited to: photosynthesis (Minagawa and Tokutsu, 2015), cell cycle (Cross and Umen, 2015), lipid accumulation (Li-Beisson et al., 2015), stress responses (Erickson et al., 2015; Schroda et al., 2015), and biofuel production (Scranton et al., 2015). Chlamydomonas has several advantages that can be leveraged: (1) Vegetative cells are haploid, therefore mutant phenotypes present easily; (2) Cells grow quickly with 6∼8 h doubling time under normal growth conditions; (3) Cells grow in light using photosynthesis but also in dark with a supplied carbon source, allowing for maintenance of photosynthetic mutants in the dark; (4) All three genomes (nucleus, chloroplast, mitochondrion) of Chlamydomonas are sequenced and transformable, making it an excellent model to study inter-organellar communications (Maul et al., 2002; Merchant et al., 2007; Gallaher et al., 2018); (5) It has generally smaller gene families than land plants, simplifying genetic and functional analyses (Merchant et al., 2007; Karpowicz et al., 2011); and (6) Many cellular processes in Chlamydomonas have similarities with land plants, making it an excellent model organism to identify novel genes/pathways with potential applications in crops.

Chlamydomonas has rich genetic and genomic resources, and its unicellular nature enables high-throughput approaches and functional genomics. Many transcriptomic analyses under various conditions exist in Chlamydomonas (Salomé and Merchant, 2021), including light stress (Mettler et al., 2014), nutrient limitation (González-Ballester et al., 2010; Schmollinger et al., 2014; Park et al., 2015), metal deficiency (Urzica et al., 2012; Blaby-Haas et al., 2016), day/night cycle (Zones et al., 2015; Strenkert et al., 2019), oxidative stress (Blaby et al., 2015), and temperature stresses (Légeret et al., 2016; Li et al., 2020; Zhang et al., 2021). Additionally, several proteomics datasets for specific cellular structures are available, including chloroplasts (Terashima et al., 2011), pyrenoids (Mackinder et al., 2016; Zhan et al., 2018), mitochondria (Atteia et al., 2009), and others. Two powerful and efficient gene cloning approaches are well-established in Chlamydomonas: the Moclo Toolkit (Golden Gate cloning kit for synthetic biology) (Crozet et al., 2018) and the recombineering pipeline to clone large and complex genes (Emrich-Mills et al., 2021). Furthermore, a genome-saturating, indexed, mapped Chlamydomonas mutant library is available for both reverse and forward genetic screens (Chlamydomonas Library Project, CLiP, https://www.chlamylibrary.org/) (Li et al., 2016, 2019). The mutant library also allows for high-throughput, quantitative phenotyping of genome-wide mutants in pooled cultures under different conditions (Li et al., 2019; Vilarrasa-Blasi et al., 2020). The mutant library is maintained and distributed by the Chlamydomonas Resource Center at the University of Minnesota, which also has several other collections of mutants in Chlamydomonas (https://www.chlamycollection.org/). Additionally, highly efficient insertional mutagenesis and CRISPR gene editing tools are well-optimized in Chlamydomonas (Shimogawara et al., 1998; Wang et al., 2019; Greiner et al., 2017; Dhokane et al., 2020; Picariello et al., 2020). These resources and tools accelerate research using Chlamydomonas as a model organism.

Two commonly used Chlamydomonas background strains are CC-1690 and CC-5325. CC in the strain names is short for <u>C</u>hlamydomonas Resource <u>C</u>enter. CC-1690 is also called *21gr* or Sager 21gr because it was from Ruth Sager at the Sidney Farber Cancer Institute in 1983 (Sager, 1955; Pröschold et al., 2005). CC-1690 is mating type plus (mt^+^) and has been used for most of the cDNA libraries and subsequent EST sequences, and the Chlamydomonas Genome Project (Shrager et al., 2003). Its genome has been sequenced multiple times by several laboratories (Gallaher et al., 2015; Flowers et al., 2015). It has also been used for several transcriptome analyses (Mettler et al., 2014; Zhang et al., 2021) and is an often-preferred background strain for photosynthesis- or stress-related experiments (Mackinder et al., 2016; Yamano et al., 2021; Zachleder et al., 2019; Ivanov et al., 2021). CC-5325 is relatively new, available to the Chlamydomonas community since 2014 (Zhang et al., 2014), but has been frequently used because it is the background strain of the Chlamydomonas Library Project (CLiP, https://www.chlamylibrary.org) (Li et al., 2016, 2019). CC-5325 is mating type minus (mt^-^) and it is the same as CC-4533, which was from an independent cryogenic storage copy. CC-5325 is also called CMJ030 (the <u>30</u>^th^ select <u>C</u>hlamydomonas strain from the laboratory of Dr. <u>M</u>artin <u>J</u>onikas). CC-5325 was isolated from a cross between two parental strains: 4A^-^ (cell wall intact) and D66^+^ (cell wall deficient). This particular strain was chosen as the background strain of the CLiP mutant library because of its several desirable features (Zhang et al., 2014; Li et al., 2016): it recovers well from cryogenic storage, has high electroporation and mating efficiencies, grows efficiently in heterotrophic and photoautotrophic conditions, has a functional carbon concentrating mechanism (CCM) and low clumping in liquid culture, swims normally, and has minimal adherence to glass. Since 2015, more than 500 scientific papers have been published using CC-5325 or mutants in its background, for example (Mackinder et al., 2016; Chaux et al., 2017; Itakura et al., 2019; Mukherjee et al., 2019; Perlaza et al., 2019; He et al., 2020; Amiya and Shapira, 2021). Genome-wide mutant screens have been performed under diverse growth conditions by using the Chlamydomonas CLiP mutant library (Li et al., 2019; Vilarrasa-Blasi et al., 2020).

Despite its widespread use, some questions regarding CC-5325 remain. Although a genome-wide screen for mutants deficient in photosynthesis was performed by using the CLiP mutant library (Li et al., 2019), detailed analysis of photosynthesis in CC-5325, the CLiP background, is rare. Chaux et al. reported some photosynthetic measurements in CC-5325 when characterizing a CLiP mutant with deficiency in the light reaction of photosynthesis (Chaux et al., 2017). However, photosynthesis in CC-5325 has not been thoroughly evaluated by comparing with other commonly used laboratory strains, such as CC-1690. This information will be helpful for employing the CLiP mutant library to study mutants deficient in photosynthesis. Additionally, because one parent strain of CC-5325, D66^+^, is cell-wall deficient, CC-5325 is often referred to as a cell-wall deficient strain and the Chlamydomonas Resource Center labels it with *cw15* for cell wall deficiency. The name of *cw15* comes from a cell-wall deficient mutant, in which the cell wall is absent or greatly reduced as compared to wildtype (Davies and Plaskitt, 1971; Hyams and Davies, 1972). Consequently, to eliminate the effects of cell wall deficiency, researchers often cross CLiP mutants in the CC-5325 background to other cell-wall intact strains for further analysis. However, the integrity of the cell wall in CC-5325 has not been carefully investigated. Furthermore, among the 121 different conditions for genome-wide Chlamydomonas mutant screens using the CLiP mutant library, screens related to circadian rhythms are lacking (Vilarrasa-Blasi et al., 2020). It is unclear if CC-5325 displays normal circadian rhythms and how well it can maintain its circadian rhythms.

To address these questions mentioned above, we performed and compared the phenotypic analyses in CC-1690 and CC-5325. Our results showed that: (1) CC-5325 had lower photosynthetic efficiency than CC-1690, but it grew better than CC-1690 in dark conditions, suggesting CC-5325 is a suitable backgrounds strain for mutants deficient in photosynthesis; (2) CC-5325 was more heat sensitive than CC-1690; (3) CC-5325 had an intact cell wall with comparable integrity to that in CC-1690, except that the cell wall in CC-5325 appeared thinner than that in CC-1690; (4) CC-1690 could maintain robust circadian rhythm of phototaxis in either constant darkness or constant light, but CC-5325 could not. In this work, we focused on specific phenotypes related to photosynthesis, thermotolerance, and circadian rhythm of phototaxis in these two strains. We believe our results will be useful for the Chlamydomonas community to better understand the differences between these two commonly used background strains, especially CC-5325 as the background of the CLiP mutant library.

## Results

We grew Chlamydomonas strains CC-1690 and CC-5325 in photobioreactors (PBRs) under well-controlled conditions. Under the same cultivation conditions in PBRs, these two strains had different cell shapes and sizes (Figure 1). CC-5325 had larger cell volume than CC-1690. Despite these morphological differences, both had comparable growth rates, as estimated by the rate of increased OD_680_ or chlorophyll content (Figure 2).

**Figure 1.**
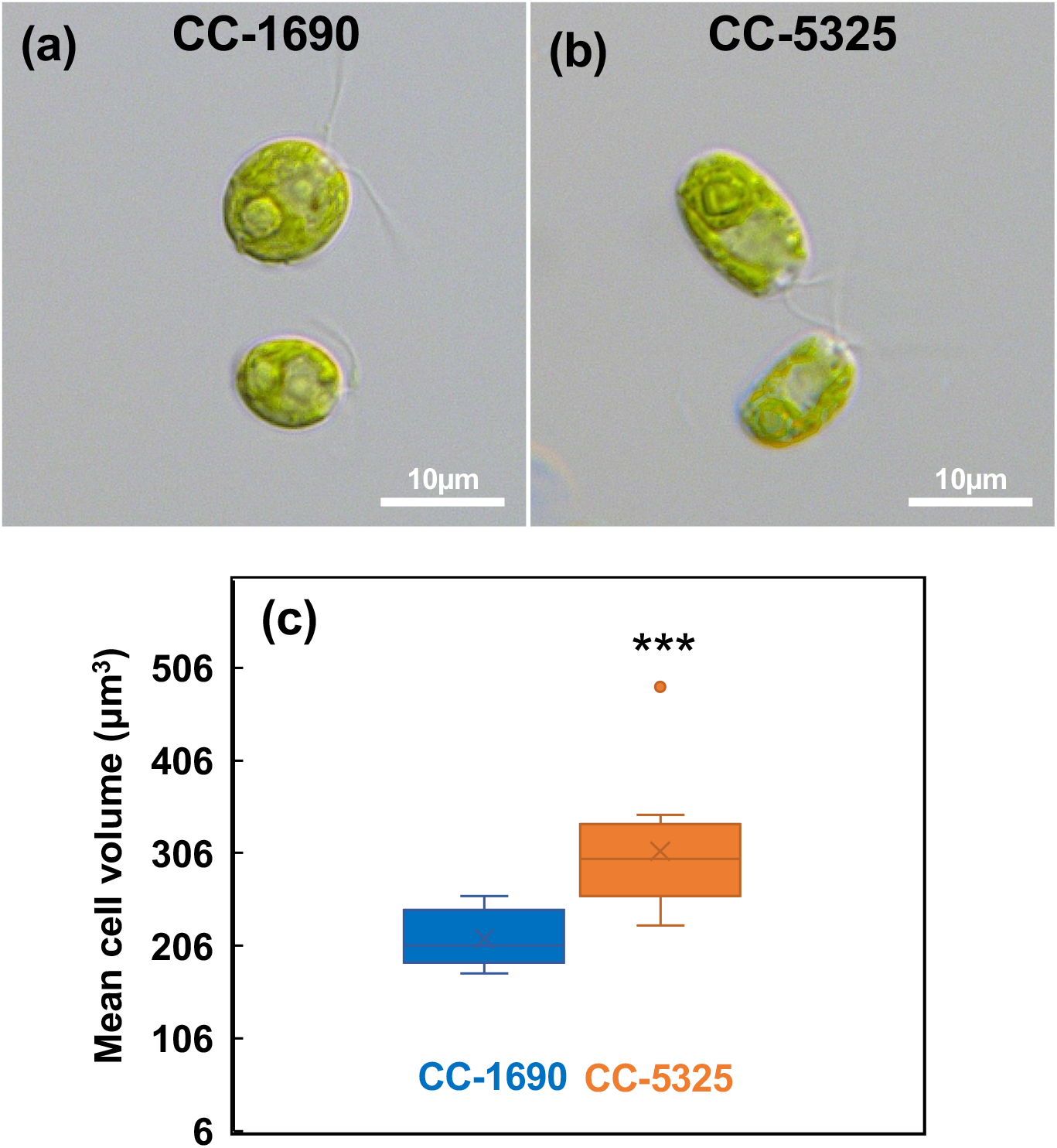
CC-5325 had larger cell volume than CC-1690. (**a, b**) Light microscopic images of Chlamydomonas cells. (**c**) Boxplot of mean cell volume determined using a Coulter Counter. n=12 biological replicates for each strain. Statistical analyses were performed using a two-tailed t-test assuming unequal variance by comparing with CC-1690. ***, P<0.001.

**Figure 2.**
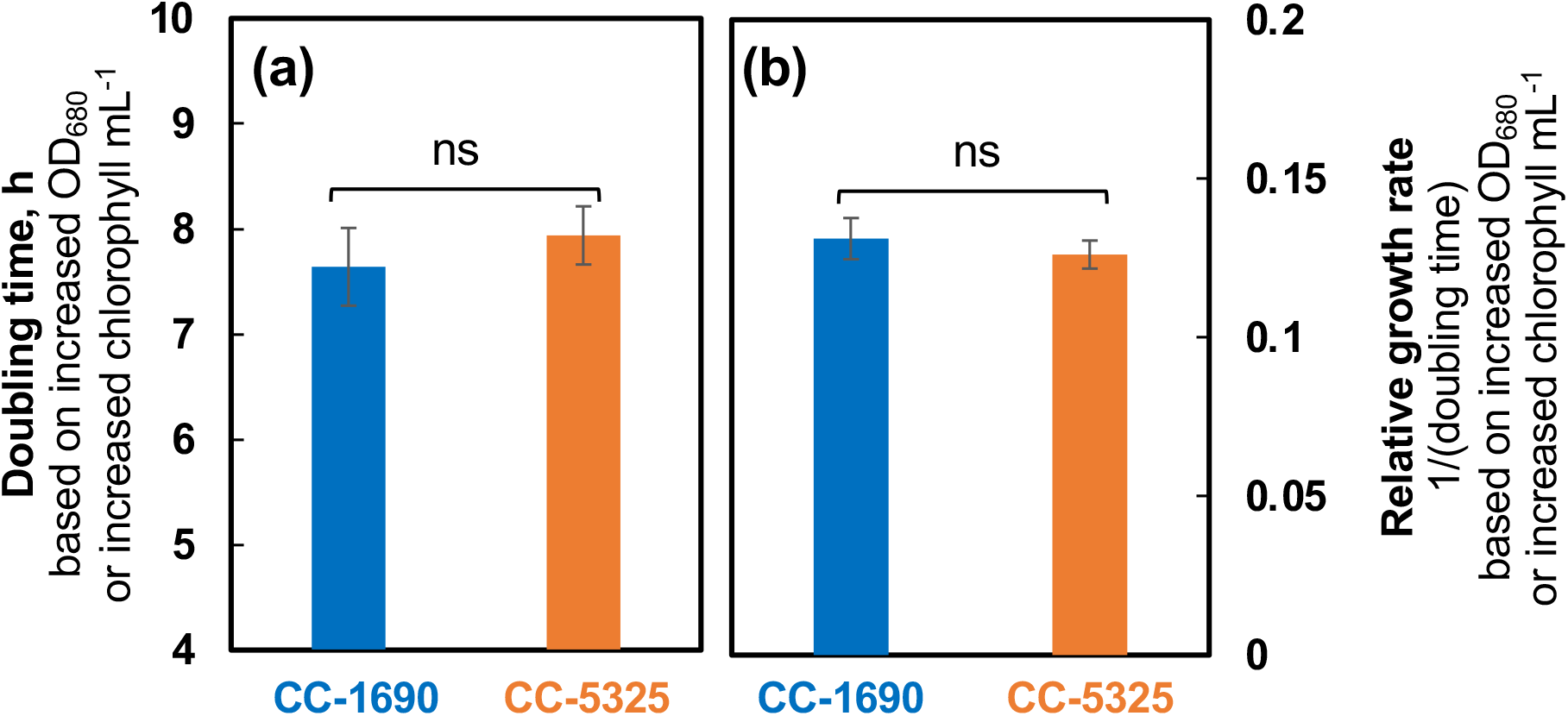
CC-1690 and CC-5325 had comparable growth rates under mixotrophic conditions in photobioreactors. Chlamydomonas cells were grown in turbidostatically controlled photobioreactors (PBRs) in Tris-acetatephosphate (TAP) medium at 25°C with a light intensity of 100 μmol photons m^-2^ s^-1^ and constantly bubbling of air. Doubling time (**a**) and relative growth rates (**b**, inverse of doubling time) were calculated based on the exponential increase of OD_680_, which is proportional to total chlorophyll content in unit of μg chlorophyll mL^-1^ and cell density at 25°C. Mean ± SE, n=3 biological replicates. Statistical analyses were performed using a two-tailed t-test assuming unequal variance by comparing the two strains; ns, not significant.

We next evaluated pigment content using algal cultures of CC-1690 and CC-5325 grown under well-controlled conditions in PBRs as mentioned above. CC-5325 had more chlorophyll and carotenoids per cell than CC-1690, but the differences were mainly due to the larger cell volume of CC-5325, as these two strains had no differences in pigments per cell volume (Figure 3a-d). Both strains had the same chlorophyll a/b ratio (Figure 3e). However, CC-1690 had a higher chlorophyll/carotenoid ratio than CC-5325, suggesting more carotenoids in CC-5325 than CC-1690 (Figure 3f).

**Figure 3.**
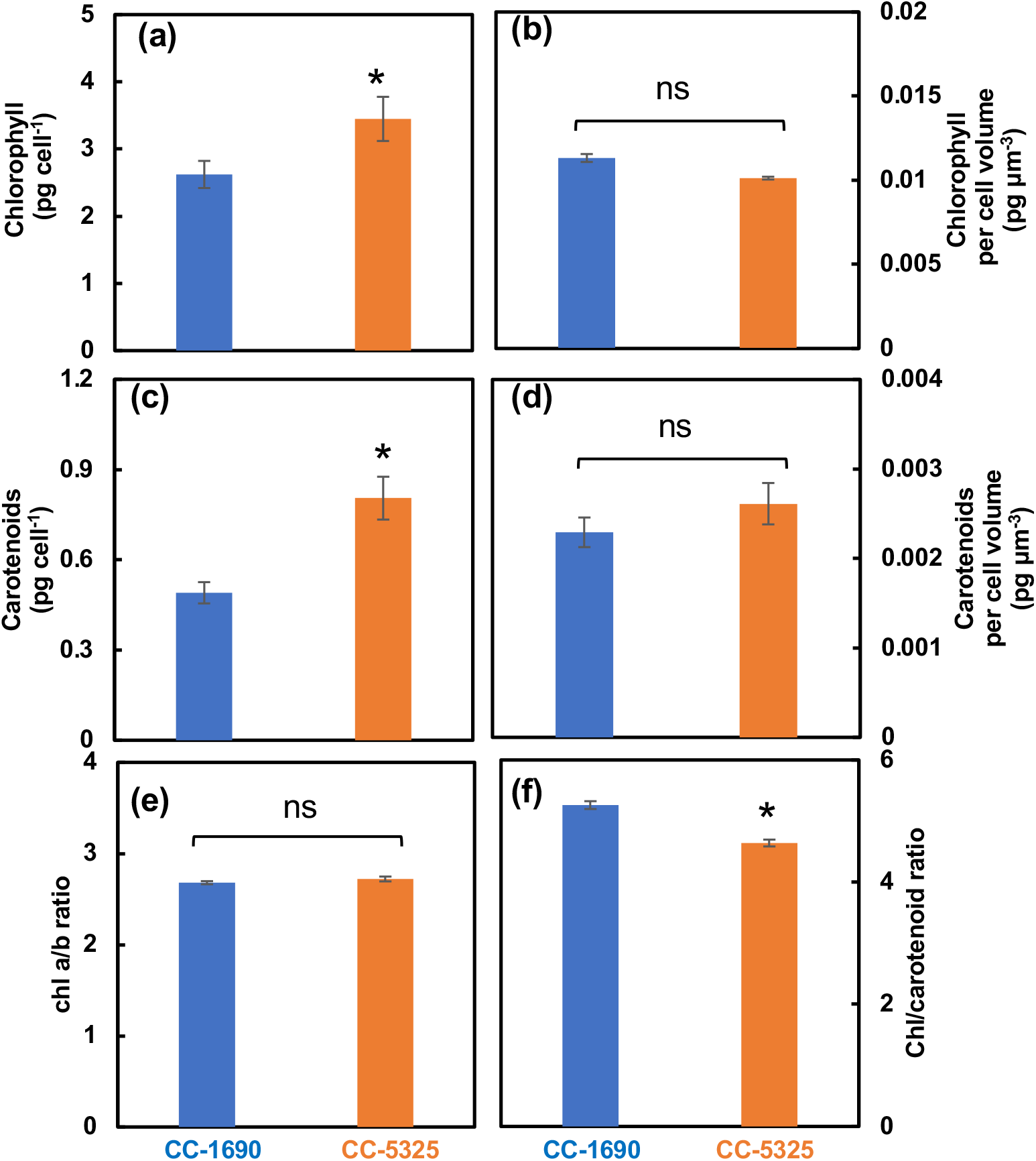
CC-5325 had more chlorophyll and carotenoids per cell than CC-1690, mainly due to the larger cell volume of CC-5325. Chlorophyll and carotenoids measured by absorbance change using a spectrophotometer. Mean ± SE, n=3 biological replicates. Statistical analyses were performed using a two-tailed t-test assuming unequal variance by comparing with CC-1690: *, P<0.05; ns, not significant.

To evaluate photosynthetic performance, we imaged room temperature chlorophyll fluorescence in algal spots grown on TAP plates using FluorCam, a chlorophyll fluorescence imager (Photon System Instruments). Both strains had indistinguishable PSII efficiency in dark and light on plates. However, CC-5325 had higher non-photochemical quenching (NPQ) than CC-1690 (Figure 4a). NPQ is one of the most important photoprotective pathways in photosynthetic organisms, helps dissipate excess light energy, and protects photosynthesis (Dietz, 2015; Müller et al., 2001). To investigate photosynthetic parameters further, we used liquid algal cultures for detailed photosynthetic measurements. We performed 77 K chlorophyll fluorescence to check the relative antenna size of PSII and PSI (Lamb et al., 2018). CC-5325 had smaller PSII antenna size than CC-1690 (Figure 4b, c). By using a custom-designed kinetic spectrophotometer/fluorometer, we showed that CC-5325 had higher PSII maximum efficiency in dark-adapted cells but lower PSII operating efficiency under 400 μmol photons m^-2^ s^-1^ light than CC-1690 (Figure 4d). In response to the same light intensities, both strains had little difference in Q_A_ redox status, except for 200 μmol photons m^-2^ s^-1^ light (Figure 4e). By using the PSII% (relative PSII antenna size or percentage of light distributed to PSII) estimated from our 77 K chlorophyll fluorescence data and PSII efficiency measured by room temperature chlorophyll fluorescence, we quantified linear electron flow (LEF) in response to light in these two strains (Figure 4f). CC-1690 had higher LEF than CC-5325 only at 400 μmol photons m^-2^ s^-1^ light. However, CC-5325 had much higher NPQ than CC-1690 under all three light intensities tested (Figure 4g). Additionally, by using a Hansatech Chlorolab 2 Clark-type oxygen electrode, we found CC-1690 had higher gross O_2_ evolution rates than CC-5325 in response to the same light intensities while both had no differences in respiration rates (Figure 4h, i). Our results suggested that CC-5325 had lower photosynthetic efficiency than CC-1690.

**Figure 4.**
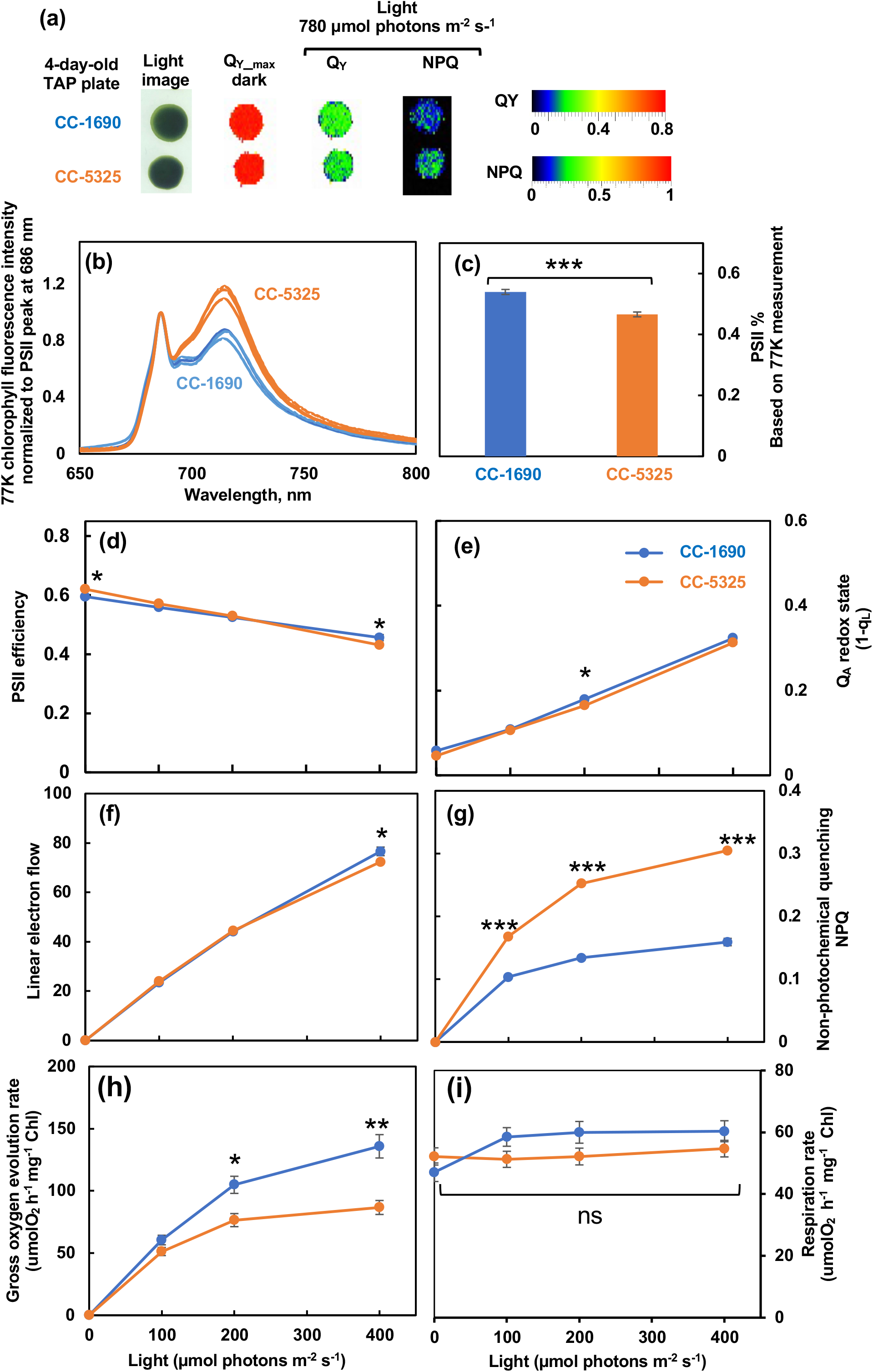
CC-1690 had higher photosynthetic efficiency than CC-5325. (**a**) Chlorophyll fluorescence imaging of 4-day-old algal spots grown on TAP plates at 25°C under 150 μmol photons m^-2^ s^-1^ white LED lights in incubators. Imaging was performed by using FluorCam (Photo System Instruments). Q_Y-max_, maximum PSII efficiency in dark-adapted plates. Under 780 μmol photons m^-2^ s^-1^ light, Q_Y_ and NPQ are PSII operating efficiency and non-photochemical quenching (NPQ) in light adapted plates, respectively. Color bars represent values of Q_Y_ and NPQ. (**b-i**) Algal cells grown in PBRs under the same condition as in Figure 2 were harvested for photosynthetic measurements. (**b**) 77 K chlorophyll fluorescence normalized to PSII peak at 686 nm. (**c**) PSII%, relative PSII antenna size, or light distributed to PSII, estimated from 77 K chlorophyll fluorescence data in (**b**). (**d-g**) Room temperature chlorophyll fluorescence measurements were performed in algal liquid cultures using a custom-designed spectrophotometer, see methods for details. (**d**) PSII efficiency, the data at 0 μmol photons m^-2^ s^-1^ light is the maximum PSII efficiency in dark and data from light phase is PSII operating efficiency in light-adapted cells. (**e**) Q_A_ redox state, the redox state of chloroplastic quinone A (Q_A_), the primary electron acceptor downstream of PSII; the bigger number of Q_A_ redox state means more reduced Q_A_. (**f**) Linear electron flow. (**g**) Non-photochemical quenching, NPQ. (**h, i**) Gross O_2_ evolution rates and respiration rates, measured using a Hansatech Chlorolab 2 Clark-type oxygen electrode. Mean± SE, n=3 biological replicates. (**c-i**) All figures have error bars, but some error bars may be too small to be visible on the graph. Statistical analyses were performed using a two-tailed t-test assuming unequal variance by comparing with CC-1690 under the same experimental conditions. *, 0.01<P<0.05; **, 0.001<P <0.01; ***, P<0.001; ns, not significant. (**d-i**) The positions of asterisks match the corresponding light intensities.

We harvested algal cultures of CC-1690 and CC-5325 from PBRs, spotted them on agar plates with either TAP (with acetate, the supplied carbon source) or TP (Tris-phosphate medium, without acetate) medium, and grew them under different light and temperature conditions in incubators (Figure 5). CC-5325 grew better than CC-1690 in dark with acetate (Figure 5a). CC-1690 could grow in dark but appeared to have retarded growth as compared to CC-5325. At 25°C and 35°C under 150 μmol photons m^-2^ s^-1^ light in incubators, CC-1690 and CC-5325 had indistinguishable growth (Figure 5b-e, g). To evaluate the thermotolerance of these two strains under well-controlled temperature conditions in liquid cultures, we exposed the algal cultures to 43°C for 2 h in PBRs, then spotted the cells on agar plates to measure viability. CC-5325 barely survived the heat of 43°C; while CC-1690 could survive 43°C heat better (Figure 5f). Our results showed that both strains grew well photoautotropically and mixotrophically, but CC-5325 grew better under heterotrophic conditions while CC-1690 had higher thermotolerance than CC-5325.

**Figure 5.**
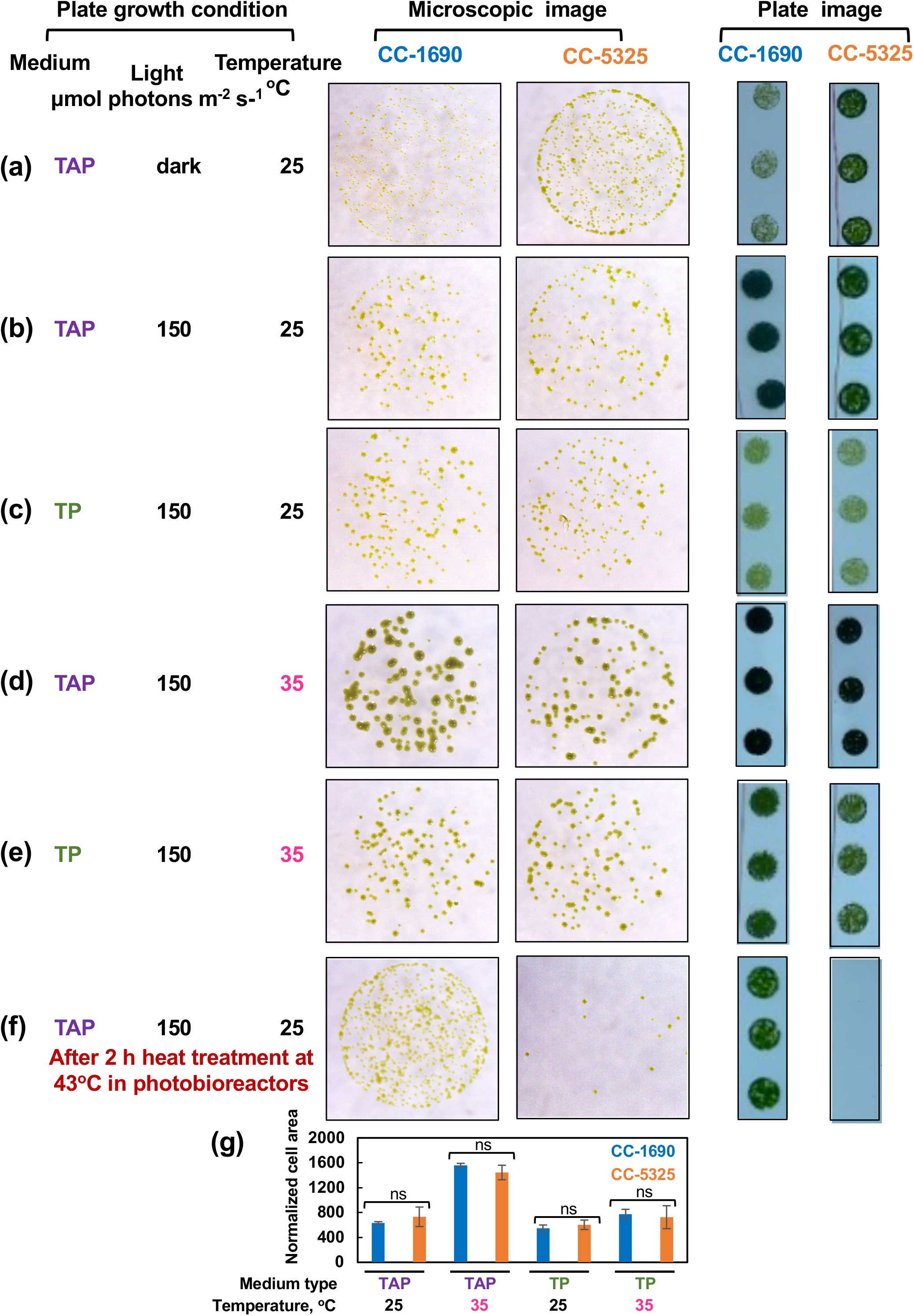
CC-5325 grew better in dark with acetate but was more heat sensitive than CC-1690. (**a-e**) Algal cells grown in PBRs under the same condition as in Figure 2 were harvested, diluted, and spotted on agar plates, and grown under the indicated growth conditions in temperature-controlled incubators. Cultures with the same cell density were used for each spot. (**f**) Algal cells were heat-treated at 43°C for 2-h in PBRs before spotting; cultures with equal volume and the same dilution were used for each spot. TAP, Tris-acetate-phosphate medium with acetate. TP, Tris-phosphate medium without acetate. The same dilution and growth duration were used for the two strains under the same condition. (**a, f**) Algal cultures with 1:20 dilution, about 1000 cells in 10 μL, were used for spotting. (**b-e**) Algal cultures with 1:100 dilution, about 200 cells in 10 μL, were used for spotting. Single algal spots were imaged using a dissect microscope after 5-day (**a**) or 44-h (**b-f**) of growth. Plate images (all 1: 20 dilution) were taken after 10-day (**a**) or 3-days (**b-f**) of growth using a regular camera. Images shown are representative of three biological replicates. (**g**) Normalized cell area. The 44-h spotting images were analyzed by ImageJ to get cell areas which were then normalized to the number of cells spotted. Statistical analyses were performed using a two-tailed t-test assuming unequal variance by comparing with CC-1690 under the same experimental conditions; ns, not significant. Such quantification was not performed for (**a, f**) because the cell areas for one of the strains were too small to do the quantification.

Thermotolerance has been linked to cell wall properties (Xu et al., 2014; Gall et al., 2015), although it is unclear how cell walls affect thermotolerance. Because CC-5325 is a progeny from a cross between 4A^-^ (cell wall intact) and D66^+^ (cell wall deficient), CC-5325 is often thought to be cell wall deficient. We evaluated the cell walls of CC-1690 and CC-5325 based on freezing tolerance, a detergent assay, and cell wall imaging (Figure 6-8). Cell-wall deficient strains are thought to recover better from liquid nitrogen freezing than cell-wall intact strains (Harris et al., 2009). Both CC-1690 and CC-5325 recovered equally well from liquid nitrogen freezing (Figure 6). Next, we used a detergent assay to evaluate the integrity of the cell wall. Triton X-100 is a detergent to disrupt lipid-based membranes and cause cells without an intact cell wall to lyse. Strains without intact cell wall or reduced cell wall integrity will have more damaged cells in the presence of Triton X-100, thus releasing more chlorophyll into the supernatant, which could be quantified using absorbance at 435 nm. We included the well-studied cell-wall-deficient line *cw15* as a control for this assay (Davies and Plaskitt, 1971; Hyams and Davies, 1972). Without Triton X-100, there was little chlorophyll released in all three lines (Figure 7a). With Triton X-100, all three lines had increased chlorophyll release, but CC-5325 had the least chlorophyll release while *cw15* had the most, demonstrating the effectiveness of our assay. By using a Coulter Counter, we quantified the number of intact cells with and without Triton X-100 (Figure 7b). While Triton X-100 did not affect CC-1690 or CC-5325, it damaged nearly all cells of *cw15*. Our freezing and detergent assays showed that CC-1690 and CC-5325 had comparable cell wall integrity.

**Figure 6.**
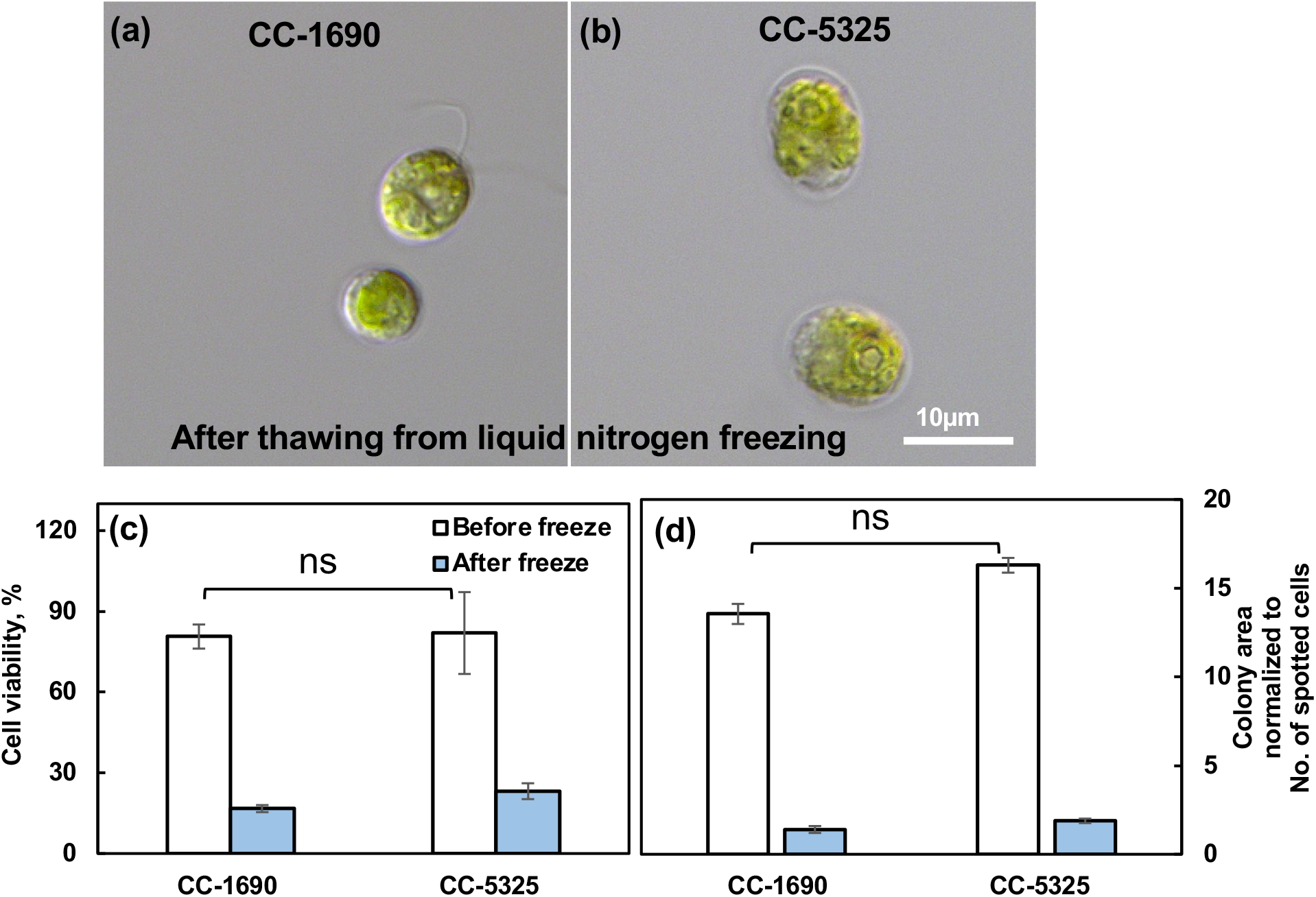
CC-1690 and CC-5325 recovered from liquid nitrogen freezing equally well. (**a, b**) Representative images of CC-1690 and CC-5325 after thawing from liquid nitrogen freezing. (**c**) Cell viability before and after freezing quantified by the No. of colonies on TAP plates. (**d**) Colony area on TAP plates for cells before and after freezing, normalized to the total number of cells spotted for each condition. The numbers of intact cells before and after freezing in liquid cultures were measured using a Coulter Counter and diluted cultures were spotted on TAP plates. TAP plates with algal spots were grown at 25°C under 150 μmol photons m^-2^ s^-1^ white LED lights for 44-h before plate imaging. Colony numbers and areas were quantified using ImageJ. Cell viability was calculated by the No. of colonies on TAP plates divided by the No. of cells spotted. Mean± SE, n=3 biological replicates. Statistical analyses were performed using a two-tailed t-test assuming unequal variance by comparing the two strains under the same experimental conditions. No significant differences between these two strains under the same conditions, ns, not significant, P>0.05.

**Figure 7.**
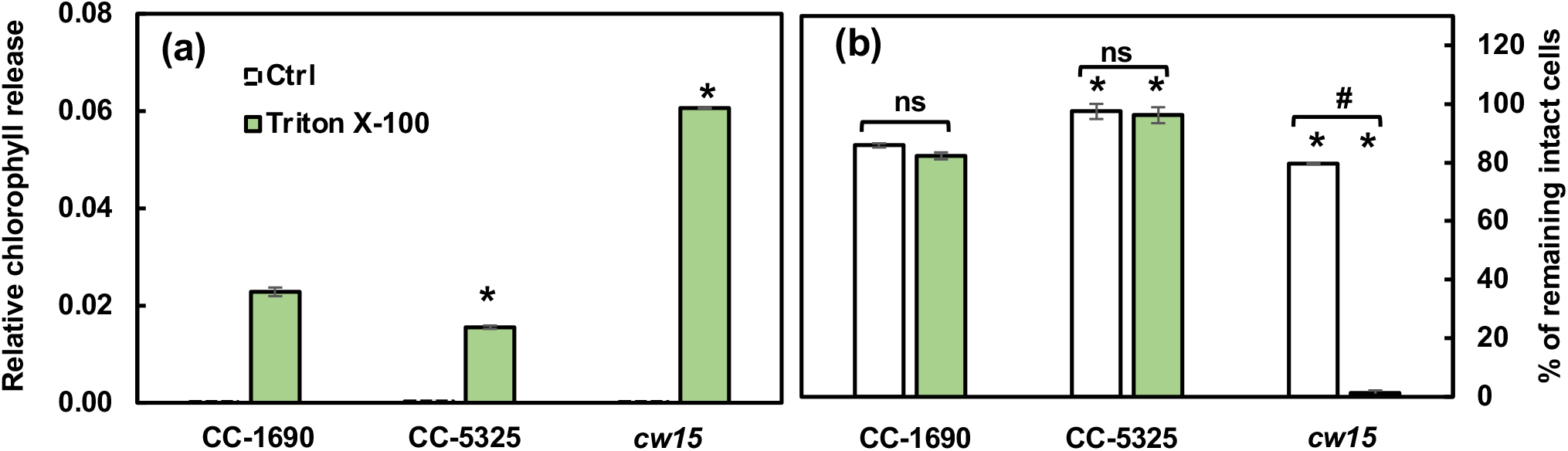
CC-1690 and CC-5325 had comparable resistance to the detergent Triton X-100. Detergent assay was performed to check cell wall integrity. Triton X-100 is a detergent to disrupt lipid-based membranes and cause cells without an intact cell wall to lyse. Algal samples with and without 0.05% Triton X-100 were vortexed vigorously. Strains without intact cell walls will have more broken cells in the presence of Triton X-100 and thus more released chlorophyll than strains with intact cell walls. (**a**) Released chlorophyll from damaged cells were quantified using absorbance at 435 nm. (**b**) The percentage of intact cells were quantified using a Coulter Counter. Mean± SE, n=3 biological replicates. Statistical analyses were performed using a two-tailed t-test assuming unequal variance by comparing with CC-1690 under the same experimental conditions (*, P<0.05) or within the same strain in the presence or absence of Triton X-100 (#, P<0.05; ns, not significant, P>0.05).

To directly visualize and quantify the cell wall, we used the lectin Concanavaline A as a cell wall stain and Syto 13™ Green Fluorescent Nucleic Acid to label the nucleus, then imaged the cells using a ZEISS Elyra 7 super-resolution microscope (Figure 8). Our results showed that CC-1690 and CC-5325 had continuous and intact cell walls, while *cw15* has discontinuous and broken cell walls, as expected for a cell-wall deficient mutant (Figure 8a-f). We calculated the total cell wall fluorescence intensity along each cell and normalized it to the cell perimeter length. The mean cell wall fluorescence intensity of CC-5325 was 84% of that in CC-1690, while that in *cw15* was 17% relative to CC-1690 (Figure 8g). Based on our results of freezing recovery, detergent assays, and cell wall imaging, we conclude that CC-5325 is not that cell-wall deficient as previously described, instead it has intact cell wall with comparable integrity as that in CC-1690, but possibly thinner cell wall than that of CC-1690.

**Figure 8.**
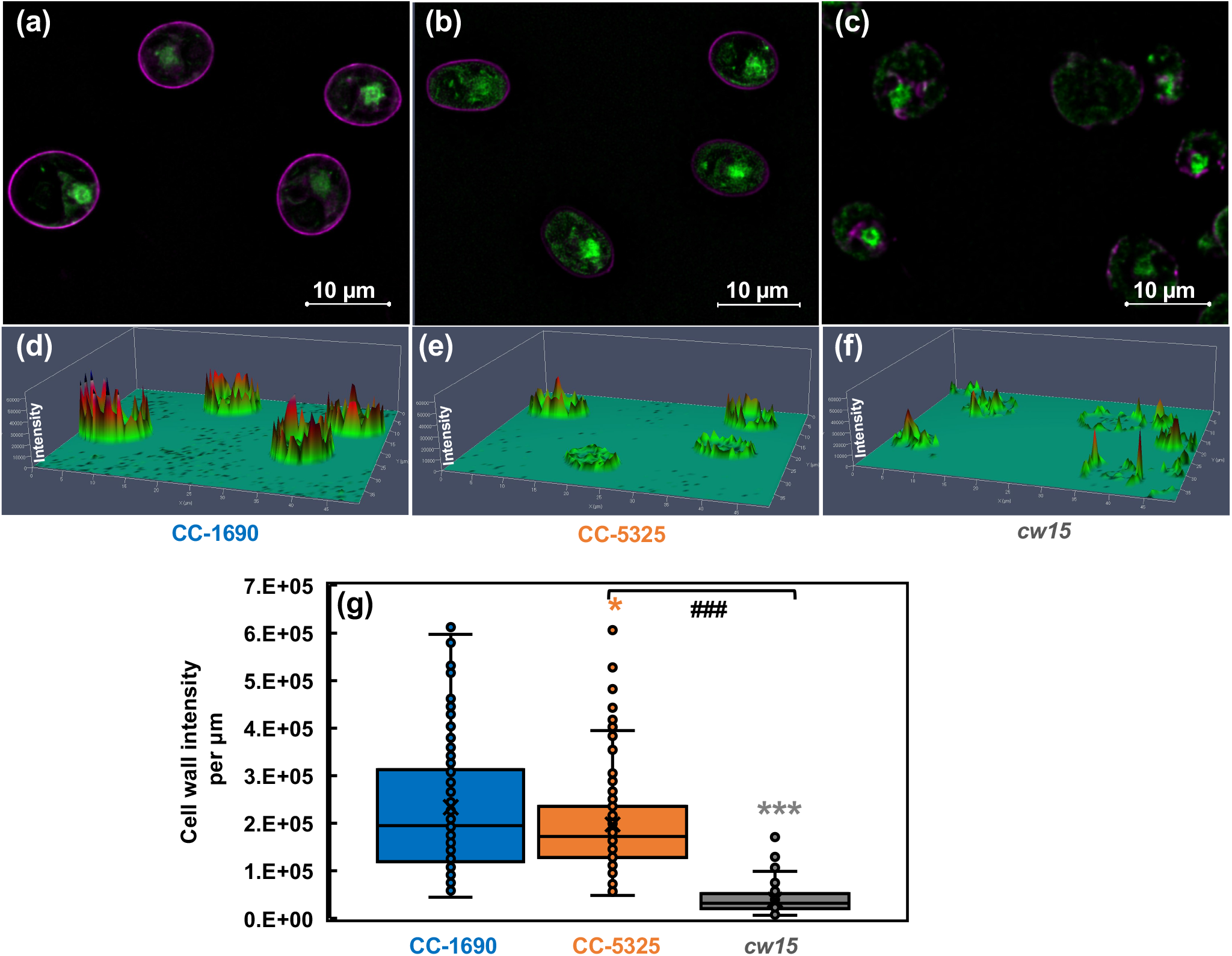
CC-5325 had intact cell wall which may be thinner than that in CC-1690. CC-1690, CC-5325 and *cw15* cells were stained with Concanavalin A for cell wall imaging and Syto 13™ Green Fluorescent Nucleic Acid to label the nucleus. *cw15*, a cell-wall-deficient line, was used as a control. (**a-c**) Representative images for cell wall stain (majenta color) and nuclear stain (green color) for three different strains. (**d-f**) Cell wall fluorescence quantification; the scale for cell wall fluorescence intensity is in the vertical direction, taller peaks represent greater fluorescence intensity. (**a**,**d**), (**b, e**), (**c**,**f**): data for CC-1690, CC-5325, and *cw15*, respectively. (**g**) Boxplot for cell wall fluorescence quantification. Total fluorescence intensity of each cell wall was normalized to the cell perimeter length. About 100 cells were quantified for each strain for fluorescence intensity. Statistical analyses were performed using a two-tailed t-test assuming unequal variance by comparing with CC-1690 (*, P<0.05; ***, P<0.001) or between CC-5325 and *cw15* (###, P<0.001).

Lastly, we investigated the circadian rhythm of phototaxis in CC-1690 and CC-5325 because both phototaxis and circadian rhythm are important for photosynthesis, thermotolerance, and cell growth (Matsuo and Ishiura, 2010; Schulze et al., 2010; Dodd et al., 2015; Blair et al., 2019). Phototaxis was determined in day/night synchronized cultures using either “constant darkness” or “constant light” free-running conditions (Figure 9). Under constant dark conditions (Figure 9a-d), the darkness was interrupted every hour by a narrow, dim light beam directed through the culture for 15 min to allow for phototaxis measurements. In constant light conditions (Figure 9e-h), the white background light was turned off every hour for the 15 min of phototaxis measurement via the narrow, dim light beam. CC-1690 showed a robust circadian rhythm of phototaxis during either constant darkness or constant light. The period of the rhythm for CC-1690 was 27.65 *±* 0.13 h in constant darkness (n=22, mean *±* se) and 23.61 *±* 0.16 h in constant light (n=12). However, in CC-5325 a sustained circadian rhythm of phototaxis could neither be detected in constant darkness nor in constant light. This inability was not simply due to an inability of the strain to undergo phototaxis (Figure 9b and f). When monitoring the phototaxis of CC-5325, there seemed to be the start of a rhythm in nearly all cultures tested in constant light and in most of the cultures tested in constant darkness but only for the first 0.5 to 1.5 days (Figure 9b and f). The lack of a sustained circadian rhythm of phototaxis in CC-5325 was true for cultures with different cell densities (from 0.5 × 10^6^ to 1.6 × 10^6^ cells mL^-1^). Our results clearly demonstrated that CC-1690 could maintain a sustained circadian rhythm of phototaxis but CC-5325 could not.

**Figure 9:**
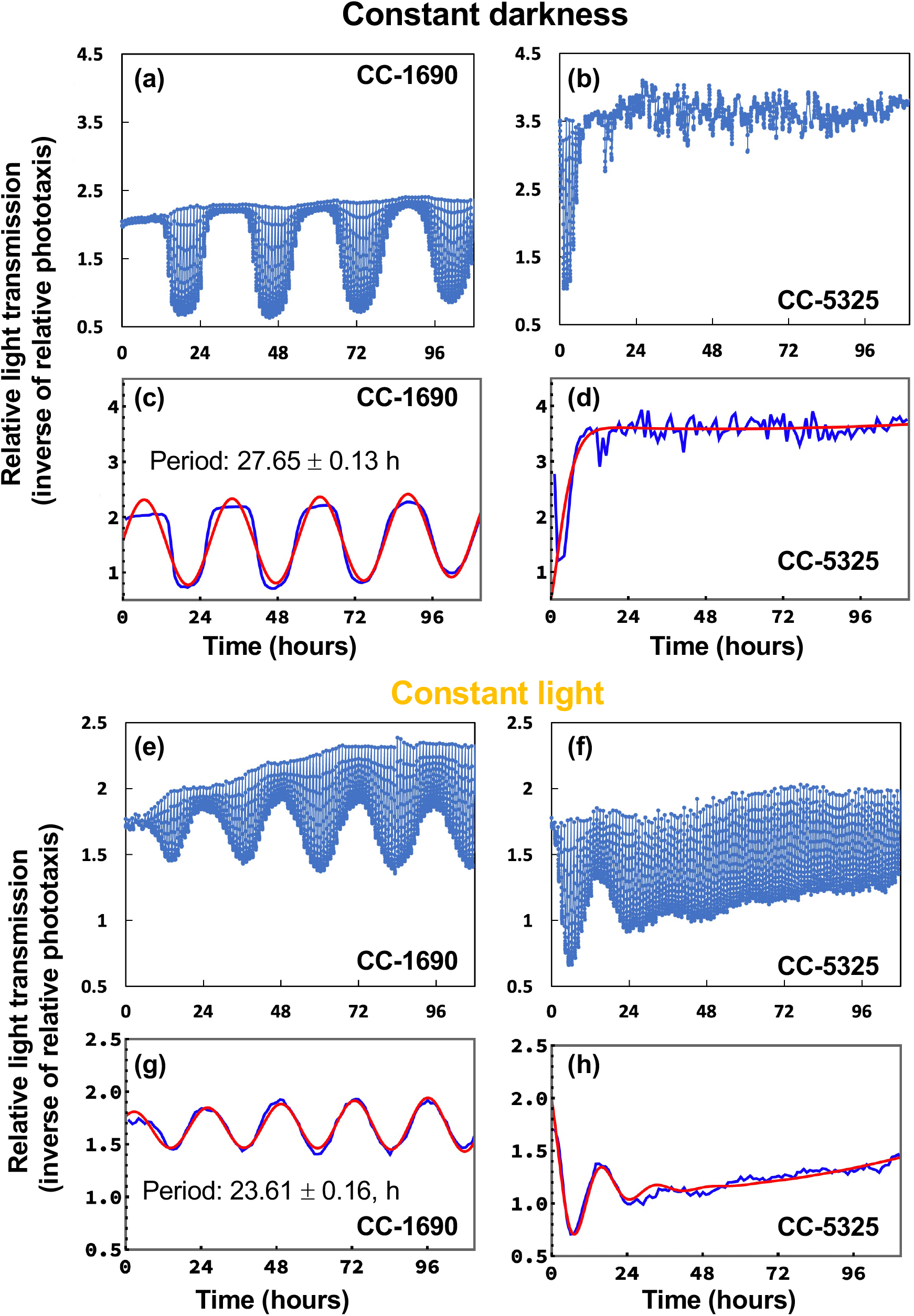
In constant darkness or constant light, CC-1690 maintained a sustained circadian rhythm of phototaxis but CC-5325 could not. All graphs on the left depict data for CC-1690 and those on the right for CC-5325. (**a, b, e, f**): Example graphs of the raw data as collected by the phototaxis machine. Each vertical line in the graph shows the light transmission measurements the machine collected at 1-minute intervals during the 15 minutes when a narrow, weak test light beam was directed through the culture sample and cells accumulated in the beam. This 15-minute test light beam was repeated every hour. During the 45 minutes that separated consecutive light beams, culture samples were exposed to constant darkness (**a-d**) or constant light (**e-h**). (**c, d, g, h**) The raw data from the above graphs (blue line) were fed into an analysis algorithm leading to the best-fit sine wave the algorithm determined for the raw data (red line). Only a single data point of the raw data was used by the algorithm per 15-minute test light cycle, the transmission measured after 11 minutes into the light beam.

## Discussion

We characterized and compared phenotypic differences between CC-1690 and CC-5325 by focusing on characteristics that are important for algal growth, including cell size, growth in liquid and on agar plates, photosynthesis, thermotolerance, cell wall integrity, and circadian rhythm of phototaxis. These two frequently used background strains displayed notable differences in these characteristics (Table 1).

**Table 1:** Summary of phenotypic differences between CC-1690 and CC-5325.

|  | CC-1690 | CC-5325 |
| --- | --- | --- |
| Other names | 21gr | CMJ030, CC-4533,<br>CLiP background |
| Cell volume |  | Larger |
| Growth in PBRs, TAP,<br>100 $\mu\text{mol photons m}^{-2} \text{ s}^{-1}$ light, air, 25°C | Similar | |
| Chlorophyll and carotenoid per cell |  | Greater |
| Chlorophyll and carotenoid per cell volume | Similar |  |
| Chlorophyll a/b ratio | Similar |  |
| Chlorophyll/carotenoid ratio |  | Smaller |
| PSII antenna size |  | Smaller |
| Non-photochemical quenching<br>NPQ |  | Increased |
| Gross O<sub>2</sub> evolution rates | | Smaller at or above 200<br>$\mu\text{mol photons m}^{-2} \text{ s}^{-1}$ light |
| Respiration rates | Similar |  |
| Heterotrophic growth in dark on plates |  | Faster |
| Mixotrophic and photoautotrophic growth<br>on plates (150 $\mu\text{mol photons m}^{-2} \text{ s}^{-1}$ light,<br>air, 25 or 35°C) | Similar | |
| Sensitivity to 43°C in PBRs |  | Greater |
| Freezing tolerance | Similar |  |
| Detergent tolerance | Similar |  |
| Cell wall integrity | Similar |  |
| Cell wall fluorescence intensity |  | Reduced |
| Circadian rhythm of phototaxis<br>in constant darkness or constant light | Yes | No |

Both strains grew well in liquid medium and on agar plates under normal conditions (Figure 2, 5). CC-5325 grew more robustly in dark with acetate than CC-1690 (Figure 5a), making it a promising background strain for mutants deficient in photosynthesis, which are often maintained in dark conditions with supplied carbon source. The reduced photosynthetic efficiency in CC-5325 may be related to smaller PSII antenna size and/or increased NPQ as compared to CC-1690 (Figure 4c, g, h). The NPQ differences in these two strains may be relative, since NPQ from different strains may not be compared directly (Maxwell and Johnson, 2000; Baker et al., 2007). The increased NPQ in CC-5325 may be related to the reduced chlorophyll/carotenoid ratio (more carotenoids) (Figure 3f). Carotenoids have reported roles in photoprotection and one of the carotenoid pigments, zeaxanthin, has major roles in NPQ formation (Baroli et al., 2004; Erickson et al., 2015). The bigger PSI peak in CC-5325 suggested its preference for state 2 and reduced PSII antenna size as compared to CC-1690 (Lamb et al., 2018). Future investigation of genes/pathways related to state transition and NPQ may help explain these differences. CC-5325 has lower oxygen evolution rates than CC-1690 only under light intensity equal to or greater 200 μmol photons m^-2^ s^-1^ light (Figure 4d), which may explain the comparable growth rates of these two strains in PBRs under 100 μmol photons m^-2^ s^-1^ light (Figure 2). Although CC-5325 had lower photosynthetic efficiency than CC-1690, it does not compromise its suitability as a background strain to screen for mutants deficient in photosynthesis. Because photosynthetic efficiencies are relative, it is not necessary for a background strain to have the highest photosynthetic efficiency to study photosynthesis mutants; normal photosynthesis would be adequate. Previous research showed that CC-5325 has a functional carbon concentrating mechanism (CCM) and several mutants deficient in CCM have been identified by using CC-5325 as a background strain (Mackinder et al., 2016; Itakura et al., 2019; Meyer et al., 2020). Mutants deficient in photosynthetic light reactions were also identified by using CLiP mutants on the background of CC-5325 (Chaux et al., 2017). Additionally, a genome-wide mutant screen has been performed to identify genes required for photosynthesis using the CLiP mutant library (Li et al., 2019). Based on our results and published literature, we believe CC-5325 is a suitable background to study photosynthesis in Chlamydomonas.

CC-5325 has been frequently referred as a cell-wall deficient line as *cw15* (Li et al., 2016; Mukherjee et al., 2019). Our results from freezing and detergent assays showed that the cell wall integrity in CC-5325 was comparable to that in CC-1690 (Figure 6 and 7). Cell wall imaging and quantification showed that CC-5325 had an intact cell wall, albeit possibly thinner or reduced cell wall than that in CC-1690 (Figure 8). The vegetative cell wall of Chlamydomonas is mainly comprised of proteins (Lee et al., 2007; Cronmiller et al., 2019). It is possible that these two strains have different quantity of cell wall proteins. There are three classifications of cell-wall-deficient mutants: (A) normal quality of cell walls that are unattached to the plasma membrane; (B) normal quality of cell walls that are attached to the plasma membrane; (C) minimal amount of cell walls (Davies and Plaskitt, 1971). The well-studied cell-wall-deficient mutant, *cw15*, belongs to classification C (Davies and Plaskitt, 1971; Cronmiller et al., 2019). Our results showed that CC-5325 was clearly not a C-type cell-wall mutant. Chlamydomonas cell walls protect cells from environmental stresses (Cronmiller et al., 2019), but the underlying mechanisms are unclear. Further investigation is needed to investigate if the reduced cell wall in CC-5325 contributes to some of its increased heat sensitivity in comparison to CC-1690 (Figure 5). Nevertheless, the cell wall integrity in CC-5325 is clearly much better than previous thought and its increased heat sensitivity as compared to CC-1690 does not affect its use for studying heat responses. The CLiP mutant library has been used to screen for heat sensitive mutants (Vilarrasa-Blasi et al., 2020). It may not be necessary to cross CLiP mutants on the background of CC-5325 to other cell-wall intact strains. Additionally, for research related to heat stress, it may generate conflicting results if the heat-sensitive CLiP mutants were crossed to other non-isogeneic lines (e.g., CC-1690) because different background strains may have different heat sensitivity, as demonstrated in our research. If crossing is needed, a possibly better option would be to cross CLiP mutants to the isogenic strain of CC-5325, which is CC-5155 (mt^+^), generated by backcrossing a mt^+^ progeny of CC-5325 (mt^-^) and CC-125 (mt^+^) to CC-5325 five times (Li et al., 2016).

The most surprising result may be the lack of a sustained circadian rhythm of phototaxis in CC-5325, although the strain can perform phototaxis (Figure 9). Our results clearly showed a sustained circadian rhythm of phototaxis in CC-1690, but CC-5325 only showed an attempt of a rhythm limited to the first day or so. This lack of a rhythm in CC-5325 was independent of whether constant light or constant darkness was present during the free-running condition. A number of different circadian rhythms have been studied in Chlamydomonas (Matsuo and Ishiura, 2010; Schulze et al., 2010), including phototaxis, chemotaxis, cell division, UV sensitivities, and others. How well Chlamydomonas can maintain these rhythms under various environmental conditions is less known. From our data, we cannot distinguish whether the lack of a sustained rhythm in CC-5325 is due to differences in its circadian clock compared to CC-1690 and therefore affecting also other circadian rhythms or whether it is specifically due to differences in how the circadian clock regulates phototaxis. If the first possibility is true, the reduced photosynthetic efficiency and thermotolerance observed in CC-5325 may be related to its deficiency in maintaining circadian rhythms, as normal circadian rhythms are important for photosynthetic organisms to optimize photosynthesis and acclimate to stressful environments (Dodd et al., 2015; García-Plazaola et al., 2017; Blair et al., 2019; Morales and Kaiser, 2020). In CC-1690, constant darkness or light affected the period of the rhythm, with a longer period in constant darkness (27.65 h) than constant light (23.61 h). Differences in period length under different constant conditions have been observed in many organisms. For example, it was recently reported that the duckweed, *Lemna gibba*, showed an average period under conditions of constant darkness that was about 7 hours longer than it was in constant light (Muranaka and Oyama, 2016). Transcriptome analysis in cultures under synchronizing versus free-running conditions may provide some insights about the mechanisms of the maintenance of circadian rhythms and about the basis for the observed differences between CC-1690 and CC-5325. Future work to investigate the underlying mechanisms by RNA-Seq or quantitative trait locus (QTL) mapping using CC-1690 and CC-5325 may help elucidate the regulation of circadian rhythms in green algae and possibly other photosynthetic organisms.

QTL mapping has been used frequently to identify causative genes related to a phenotype in land plants (Xu et al., 2017; Camargo et al., 2018; Wu et al., 2018; Wang et al., 2018). Such analysis is much less developed in Chlamydomonas. However, Chlamydomonas has several advantages for QTL mapping as compared to land plants, e.g., high mating efficiency and easy mating procedure (Jiang and Stern, 2009), faster growth on plates or in liquid, low cost in whole genome sequencing (Dutcher et al., 2012; Wakao et al., 2021), and abundant RNA-Seq datasets (Salomé and Merchant, 2021). One round of mating in Chlamydomonas only takes about 2-3 weeks, from the mating process to identifiable progenies. Due to its unicellular nature, high throughput phenotyping tools could be applied efficiently to characterize progenies of Chlamydomonas, e.g., whole plate chlorophyll fluorescence imaging (Holub et al., 2007; Massoz et al., 2017; Rühle et al., 2018), growth quantification on plates using ImageJ (Zhang et al., 2021), and circadian rhythm monitoring using an automated phototaxis machine (Gaskill et al., 2010). The Chlamydomonas CLiP mutant library allows for both forward and reserve genetic screens and candidate genes identified by QTL in Chlamydomonas can be validated using available mutants in the CLiP mutant library (Li et al., 2016). If mutants deficient in some specific genes are unavailable in the CLiP mutant library or other mutant collections at the Chlamydomonas Resource Center, mutants disrupted in genes of interest could be generated by CRISPR-Cas 9 approaches with high efficiency (Greiner et al., 2017; Dhokane et al., 2020; Picariello et al., 2020). Comparison between genome sequences of CC-1690 and CC-5325 showed that these two strains had 117059 small variants and 355 high impact small variants (Li et al., 2016), providing potential candidates for phenotyping differences. Recently, research has been conducted in Chlamydomonas to phenotype and select progenies from a cross between two Chlamydomonas parental strains under different environmental conditions and revealed possible selection-enriched genomic loci for improved stress adaptation and photosynthetic efficiency (Lucker et al., 2021), paving the way for QTL mapping in Chlamydomonas.

## Conclusions

We presented several distinct phenotypes between two commonly used laboratory background Chlamydomonas strains: CC-1690 (*21gr*) and CC-5325 (CLiP library background). Although CC-5325 has lower photosynthetic efficiency and thermotolerance than CC-1690, it does not compromise the use of CC-5325 and the CLiP mutant library to study and screen for mutants deficient in photosynthesis or heat responses. Our results showed CC-5325 had an intact cell wall with comparable integrity as that of CC-1690, which will help reduce researchers’ concerns or confusion about its cell wall deficiency. The lack of a sustained circadian rhythm of phototaxis in CC-5325 does limit the use of the CLiP library to screen for mutants deficient in circadian rhythms of phototaxis. Finally, the rich genomic and genetic resources in Chlamydomonas will enable efficient identification of responsible genes for phenotypes related to cell size, photosynthesis, thermotolerance, and circadian rhythms in the near future.

## Materials and Methods

### Strains and culture conditions

Chlamydomonas wildtype strain CC-1690 (also called *21gr*, mt^+^) and CC-5325 (also called CMJ030, identical to CC-4533, mt^-^, CLiP library background) were ordered from the Chlamydomonas resource center. Algal cultures of these two strains were maintained in photobioreactors (PBRs) as described before (Zhang et al., 2021). The normal growth condition in PBRs was: 25°C, Tris-acetate-phosphate (TAP) medium, constant 100 μmol photons m^-2^ s^-1^ light (50% red: 50% blue), constant bubbling with filtered air (0.04% CO_2_), and turbidostatic control by monitoring OD_680_ to supply fresh medium frequently and maintain cell density around 2×10^6^ cells mL^-1^. Under the normal growth condition at 25°C, OD_680_ is linearly proportional to both chlorophyll content and cell density. Fresh medium was added to the culture automatically by a peristaltic pump when the OD_680_ reached the defined maximum value to dilute the culture; the pump was stopped when the OD_680_ dropped to the defined minimum value. Algal cultures then grew up at approximately exponential rate to the defined maximum OD_680_ value before the next dilution cycle. Our OD_680_ range was sufficiently small that we expected minimal nutrient limitation during our experiment. Thus, the turbidostatic mode precisely controlled the growth condition. We calculated the doubling time or relative growth rates based on the exponential growth phase when the peristaltic pump was off. The doubling time and relative growth rates we referred to were based on the increase of OD_680_ or the total chlorophyll per mL culture. The growth rates measured by OD_680_ were consistent with medium consumption rates and cell number increase quantified by a cell counter. For the 43°C heat treatment in PBRs, the PBR temperature was gradually increased from 25°C to 43°C within 30 min, and then the PBR temperature was maintained at 43°C for 2 h. By the end of 2-h heating at 43°C, algal cultures were harvested for spotting test to evaluate viability.

### Algal spotting test

Algal spotting test was performed as described before with minor modifications (Zhang et al., 2021). CC-1690 and CC-5325 cultures harvested from PBRs were diluted to 1×10^5^ cells mL^-1^ or 0.2×10^5^ cells mL^-1^ (1:20 or 1:100 dilution) and 10 μL aliquots of the diluted cultures (about 1000 or 200 cells) were spotted on 1.5% TAP agar plates and grown in temperature-controlled incubators under the indicated conditions. For conditions with light, 150 μmol photons m^-2^ s^-1^ light light was provided by white LED lights with indicated intensity. After 44-h growth under light in incubators, algal spots were imaged by a dissecting Leica microscope. The whole plates with algal spots were imaged again using a regular camera after 3-day-growth for visual representations. Due to the slow growth in darkness, the algal spots and the whole plates were images after 5-day and 10-day, respectively.

### Pigment measurements

Pigments were quantified as described before (Zhang et al., 2021). Three biological replicates of 1 mL of PBR cultures (around 2×10^6^ cells mL^-1^) grown under normal condition of 25°C were harvested, mixed with 2.5 μL of 2% Tween20 (Sigma, P9416-100ML) to help cell pelleting, centrifuged at 18,407 g at 4°C to remove supernatant. Cell pellets were resuspended in 1 mL of HPLC grade methanol (100%, Sigma, 34860-4L-R), vortexed for 1 min, incubated in the dark for 5 min at 4°C, and centrifuged at 15,000 g at 4°C for 5 min. Top supernatant containing pigments was analyzed at 470, 652, 665 nm in a spectrophotometer (IMPLEN Nonophotometer P300) for carotenoids and chlorophyll a/b concentrations in μg mL^-1^ using the following equations: Chl a + Chl b = 22.12*A_652_ + 2.71*A_665_, Chl a = 16.29*A_665_– 8.54*A_652_, and Chl b = 30.66*A_652_ – 13.58*A_665_ (Porra et al., 1989), and carotenoids = (1000*A_470_ – 2.86*Chl a – 129.2*Chl b)/221(Wellburn, 1994).

### Photosynthetic measurements

CC-1690 and CC-5325 cultures grown at 25°C in PBRs as mentioned above were used for all photosynthetic measurements. Algal cultures were spotted on TAP plates and grown under 150 μmol photons m^-2^ s^-1^ light at 25°C in incubators for 4-day. Algal plates were dark-adapted for 20 min before chlorophyll fluorescence imaging at room temperature using FluorCam (Photo System Instruments). Photosynthetic measurements in liquid algal cultures were performed as described before (Zhang et al., 2021). 77 K chlorophyll fluorescence measurement were performed using an Ocean Optics spectrometer (cat. No. OCEAN-HDX-XR) and 430 nm excitation provided by light emitting diodes (LEDs). Spectral data were normalized to the PSII spectral maximum value at 686 nm and relative percentage of light distributed to PSII (or relative PSII antenna size, PSII%) was calculated using this formula: PSII%= normalized PSII peak / (normalized PSII peak + normalized PSI peak at 714 nm). Normalized PSII peak equals one. Room temperature chlorophyll fluorescence in liquid algal cultures were performed using a multi-wavelength kinetic spectrophotometer or fluorometer (measuring beam with 505 nm peak emission, measuring pulses of 100 μs duration) (Zhang et al., 2021). Aliquots of 2.5 mL algal cultures (around 12 μg chlorophyll) were sampled from PBRs, supplemented with 25 μL of fresh 0.5 M NaHCO_3_ in a fluorometry cuvette (C0918, Sigma-Aldrich) with constant stirring, and dark adapted with a 10-min exposure to far-red light (peak emission of 730 nm at ∼35 μmol photons m^-2^ s^-1^) before chlorophyll fluorescence measurement in dark, and subsequent actinic light phases of 100, 200, 400 μmol photons m^-2^ s^-1^ light. The actinic light was provided by LED lights with maximal emission at 620 nm. Each light phase was about 7.5 min long. A saturating flash was applied at the end of each phase to get maximum chlorophyll fluorescence (F_m_ in dark or F_m_’ in light). Photosynthetic parameters were calculated as described (Baker et al., 2007). PSII efficiency (ΦPSII) was calculated as 1-F_o_/F_m_ and 1-F’/F_m_’ for dark-adapted and light-adapted algal cells, respectively. F_o_ and F_m_ are minimal and maximal chlorophyll fluorescence in dark-adapted algal cells. F’ and F_m_’ are steady state and maximal chlorophyll fluorescence in light-adapted algal cells. Q_A_ redox status was calculated as 1-q_L_=1-(F_q_′/F_v_′)x(F_o_′/F′)=1-[(F_m_’-F’)/(F_m_’-F_o_’)]x(F_o_′/F′). q_L_ is the fraction of open PSII centres and F_q_’ is the photochemical quenching of fluorescence (Baker et al., 2007). Linear electron flow (LEF) was calculated using the formula: LEF = (actinic light) x (Qabs_fs_) x (PSII%) x (ΦPSII). *Qabs*_fs_ is fraction of absorbed light, assuming 0.8. PSII% was calculated from 77 K chlorophyll fluorescence as mentioned above. Non-photochemical quenching, NPQ, was calculated as (F_m_-F_m_’)/F_m_’. Oxygen evolution was measured using the Hansatech Chlorolab 2 based on a Clark-type oxygen electrode at room temperature as described (Zhang et al., 2021). Two-mL of cells (around 10 μg chlorophyll) supplemented with 20 μL of 0.5 M NaHCO_3_ were incubated in the dark for 10 min with stirring before the light phases of 100, 200, 400 μmol photons m^-2^ s^-1^ light. Each light lasted 5 min followed by 2 min dark. The rates of oxygen evolution and respiration were measured at the end of each light and dark phase, respectively.

### Algal cryogenic freezing and thawing

CC-1690 and CC-5325 cultures around a cell density of 2×10^6^ cells mL^-1^ were harvested from PBRs grown at 25°C and frozen in 10% cryoprotective agent (CPA, 1:10 volume ratio of 100% methanol and fresh TAP medium). Algal cultures of 450 μL was added to 450 μL of 10% CPA (final CPA concentration was 5%) in cryo-tubes, mixed gently, transferred to a CoolCell foam freezer container, incubated in a −80°C freezer for 4 h, and finally stored in a liquid nitrogen freezer. During the thawing process, cryo-tubes with algal samples were thawed in a 37°C water bath for 5 min and centrifuged at 1000 g for 5 min at room temperature to remove the supernatant. Fresh TAP medium of 900 μL was added to resuspend cell pellets, followed by dark storage of the cryo-tubes without agitation at room temperature overnight. The next day, the cryo-tubes were centrifuged again at 1000 g for 5 min at room temperature to remove the supernatant. Cell pellets were resuspended with 200 μL of fresh TAP medium and 10 μL of the resuspended algal cultures were used for the spotting test to evaluate viability as mentioned above.

### Detergent assay for cell wall integrity

CC-1690 and CC-5325 cultures around a cell density of 2×10^6^ cells mL^-1^ were harvested from PBRs grown at 25°C as mentioned above. A cell-wall deficient strain, *cw15* (Davies and Plaskitt, 1971), was received from the Umen laboratory and used as a control. *cw15* could not grow well in PBRs probably because its cell wall deficiency made it sensitive to the constant air bubbling in PBRs. So shaker cultures of *cw15* around the similar cell density were used for the detergent assay. Growth condition of *cw15* on the shaker was 25°C, around 100 μmol photons m^-2^ s^-1^ light, and in TAP medium. One-mL algal cultures were used for chlorophyll quantification (Porra et al., 1989) and another 1-mL cultures (with 0.05% Triton X-100 or with TAP medium) were immediately vortexed vigorously for 30 s then incubated in dark for 10 min. Then, 500 μL of each sample was withdrawn and centrifuged at 13,000 × g to remove cell debris. The absorbance of the supernatant was measured at 435 nm and normalized to chlorophyll contents. The number of intact cells were measured using a Coulter Counter (Multisizer 3, Beckman Counter, Brea, CA).

### Cell wall staining and quantification

Ten mL cultures of CC-1690, CC-5325 and *cw15* cells as mentioned above were collected and concentrated to 2×10^7^ cells mL^-1^ by centrifugation at 1500 g for 5 min at room temperature. Cells were stained with Concanavaline A (ConA, Alexa Fluor®594 conjugate) (100 μg mL^-1^, ThermoFisher Scientific) (Bensalem et al., 2018) and Syto 13 Green Fluorescent Nucleic Acid Stain (5 μM mL^-1^, ThermoFisher Scientific) for 30 min at room temperature. Stained cells were washed first and then fixed in 4% paraformaldehyde (EM Science, Hatfield, PA, USA) and followed by 1X rinse in TAP medium. Fluorescence images were acquired using a 40x C-Aprochromat objective lens (numerical aperture 1.2) in apotome mode with a ZEISS Elyra 7 super-resolution microscope (ZEISS, Oberkochen, Germany). The 488 nm and 561 nm laser lines with BP 420-480 + 495-550 and BP 570-620 + LP 655 emission filters were used for Syto 13 and ConA, respectively in fast frame sequential mode at 50 ms exposure using dual pco.edge 4.2 sCMOS detectors (PCO AG, Kelheim, Germany). All two-dimensional (2D) images (with 5 phases) were acquired at 49 nm x-y pixel resolution and reconstructed using the SIM^2^ module and 2D+ processing with standard live sharpness in ZEN Black 3.0 SR FP2 software. For each strain, 100 cells were quantified by free hand drawing a line intensity profile along the cell wall. The fluorescence intensity of each cell wall periphery was normalized by its perimeter length.

### Circadian rhythm of phototaxis

CC-1690 and CC-5325 were grown in 125-mL Erlenmeyer flasks containing 50-mL photoautotrophic Tris-phosphate (TP) minimum medium without acetate. Cultures were grown on a shaker at 17 to 19°C under synchronizing 12 h light/12 h dark cycles. The light intensity during the light phase was about 20 μmol photons m^-2^ s^-1^. When cultures reached cell concentrations between 3.5 × 10^5^ and 2.3 × 10^6^ cells mL^-1^, culture samples were placed into small Petri dishes and their phototaxis was monitored at 20°C as described (Gaskill et al., 2010) with the following modifications. A box around the phototaxis machine with a heating element and ventilators allowed the external temperature around the machine to be kept at a constant but slightly elevated temperature of 24°C compared to the machine. Additionally, temperature recorders (Tempo Disc, BlueMaestro) that fit into the Petri dish slots allowed for a more precise monitoring of the temperature the culture samples were exposed to. An algorithm as described before was used to analyze the phototaxis data for their periods (Gaskill et al., 2010). Three independent experiments were conducted for each of the constant darkness and constant light conditions with 1 or 2 different cultures per strain per experiment and 3 to 4 replicate samples per culture.

### Statistical analysis

A two-tailed t-test with unequal variance was used for statistical analysis. P>0.05, not significant, or ns; *, 0.01<P<0.05; **, 0.001<P <0.01; ***, P<0.001.

## Author contributions

R.Z. supervised the project. R.Z. wrote the manuscript with the help from S. J., N.Z., and K.J.C.. N.Z. performed freezing test and detergent assay. L.P. performed room temperature chlorophyll fluorescence measurements and O_2_ evolution in liquid algal cultures. H.N. initiated the project, performed FluorCam imaging, and 77 K chlorophyll fluorescence measurements. S. J. performed the experiments of circadian rhythm of phototaxis. M. X. maintained algal cultures in photobioreactors, performed spotting and thermotolerance tests. N.Z., H.N., M.X. quantified pigments. N.Z., A. K., K.J.C. performed cell wall imaging. R.Z. and M.X. polished most of the figures. R.Z., S. J., K.J.C., L.P., and H.N. helped revise the manuscript.

## Acknowledgements

This research was supported by the start-up funding from the Donald Danforth Plant Science Center (DDPSC), the Department of Energy (DOE) Basic Energy Sciences (BES) Photosynthetic Systems (PS) grant (Award No. 0019464), and the DOE Biological & Environmental Research (BER) grant (Award No. 0020400) to R.Z. We acknowledge imaging support from the Advanced Bioimaging Laboratory (RRID:SCR_018951) at the Danforth Plant Science Center and usage of the ZEISS Elyra 7 Super-Resolution Microscope acquired through an NSF Major Research Instrumentation grant (DBI-2018962). We would like to thank Dr. Dmitri Nusinow, Erin Mattoon, and Cooper Hostetler for providing feedback for the manuscript. Dr. Dmitri Nusinow and Maria Sorkin are acknowledged for initial analysis of circadian rhythms in algal cells. We also want to thank Catherine Bailey for help with some of the pigment analysis.

## Data Availability Statement

The data presented in this study are available in this article.

